# FANGORN: A quality-checked and publicly available database of full-length 16S-ITS-23S rRNA operon sequences

**DOI:** 10.1101/2022.10.04.509801

**Authors:** Calum J. Walsh, Meghana Srinivas, Douwe van Sinderen, Paul D. Cotter, John G. Kenny

## Abstract

Sequence comparison of 16S rRNA PCR amplicons is an established approach to taxonomically identify bacterial isolates and profile complex microbial communities. One potential application of recent advances in long-read sequencing technologies is to sequence entire rRNA operons and capture significantly more phylogenetic information than sequencing of the 16S rRNA (or regions thereof) alone, with the potential to increase the proportion of amplicons that can be reliably classified to lower taxonomic ranks. Here we describe *FANGORN* (**F**ull-length **A**mplicons for the **N**ext **G**eneration **O**f r**RN**a analysis), a publicly available database of quality-checked 16S-ITS-23S rRNA operons, accompanied by multiple taxonomic classifications. *FANGORN* will aid researchers in analysis of their data and act as a standardised database to allow comparison of results between studies.

## INTRODUCTION

Ribosomal RNA sequencing is an important tool for the identification and phylogenetic analysis of prokaryotes. Sanger sequencing of the entire ~1.5kbp 16S rRNA gene is commonly used to identify cultured isolates and can provide species-level resolution in most instances. However, the 16S rRNA genes of some highly-related species, such as members of the *Streptococcus mitis* group or the *Escherichia coli* and *Shigella* spp., share >99% sequence identity and can therefore not be reliably distinguished. Sequencing the longer 23S rRNA gene instead of, or in tandem with, the 16S rRNA gene can provide greater phylogenetic resolution and allow reliable separation of these challenging species. Furthermore, although Sanger sequencing is ideal for generating long, high fidelity reads, it is inefficient for profiling complex communities and so high-throughput Illumina sequencing of single hypervariable regions is used instead. This approach can generate massive quantities of highly-accurate reads but this increased quantity comes at the expense of phylogenetic resolution due to shorter read lengths. The 16S rRNA gene has also been the target of choice for metagenetics or barcoding studies due to its mix of alternating highly-conserved and hypervariable regions.

The latest generation of long-read sequencing technologies, spearheaded by Pacific Biosciences and Oxford Nanopore Technologies, has the potential to combine the benefits of sequencing long stretches of DNA in a relatively high-throughput, culture-independent, manner. Also, due to the length of reads that can be generated on these platforms, it is possible to sequence both the 16S and 23S genes on a single stretch of DNA while also capturing the internal transcribed spacer (ITS) region. This allows a greater proportion of amplicon sequences to be assigned to species and even strain-level compared to 16S rRNA sequencing [1]. Recent studies have applied this approach to profiling microbial communities [2, 3], though they had to rely on custom-built or commercial databases, which do not allow the direct comparison of results between studies. Recently, others have identified the requirement for a regularly updated and publicly available database to format the RRN structure for metagenetics analysis [4].

Here we present *FANGORN* (**F**ull-length **A**mplicons for the **N**ext **G**eneration **O**f r**RN**a analysis), an open access quality-checked database of full-length 16S-ITS-23S rRNA sequences, accompanied by taxonomic and contextual data, to act as a database to standardise analysis between studies in the same way SILVA [5], RDP [6], and Greengenes [7] function for single rRNA genes.

## DATABASE CONSTRUCTION

Two datasets were used as the basis for the *Fangorn* database, differing primarily by the taxonomy systems employed.

The first of these, referred to as *RefSeq* throughout this manuscript, was constructed from all 253,840 RefSeq genome assemblies marked ‘Latest’ on 14/07/2022. The second, referred to as *GTDB,* was constructed from all 317,541 genomes included in the most recent release (07-RS207) of the GTDB database [8].

The pipeline described below, and illustrated in Figure 1, was used to first construct databases of quality-checked 16S-ITS-23S rRNA operon (*rrn*) sequences from complete genomes and then expand this with sequences from incomplete assemblies, thereby capturing as much diversity as possible while placing a premium on genome assembly quality.

**Figure 1.**
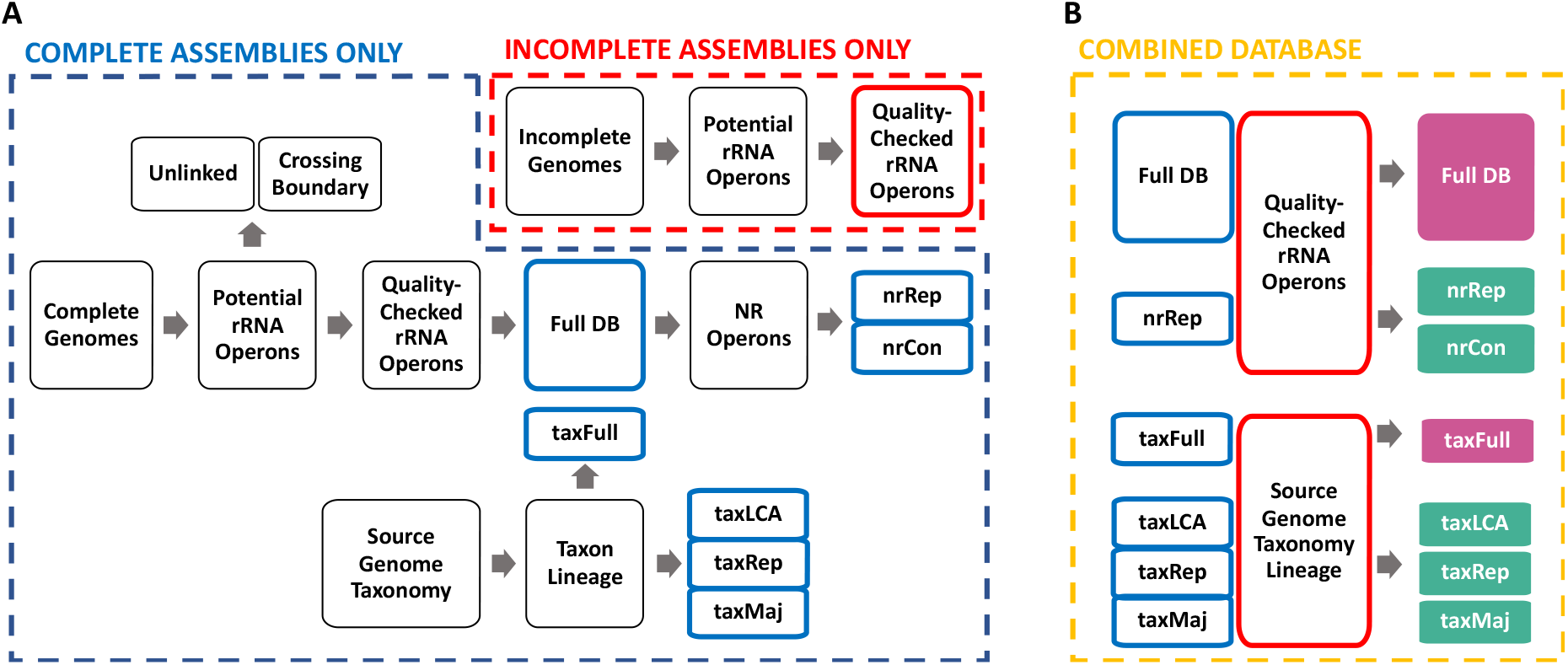
FANGORN database construction pipeline. Panel A depicts intermediary steps and files which are combined and/or dereplicated to generate the final database depicted in Panel B (more details in DATABASE CONSTRUCTION section). Colour-outlined rectangles represent intermediary files used to build the colour-filled sequence and taxonomy files available for download.

The associated annotation information of *RefSeq* and *GTDB* source genomes was downloaded in GFF3 format and rRNA gene features were extracted. If annotations were not available, rRNA genes were identified using barrnap (v 0.9) with default parameters except genes with a coverage of < 80% were marked “partial”.

The combined annotation information from complete assemblies was imported to R [9] using the read.gff function from the ape package [10], partial genes were discarded, and *rrn* sequences were identified by iteratively associating 16S genes with their neighbouring 23S gene if it was:

1. Located on the same assembled sequence (contig or scaffold)
2. Encoded on the same strand
3. Encoded in the order 16S-ITS-23S

The potential *rrn* operons identified based on these criteria were further filtered to remove ‘unlinked’ operons with ITS regions longer than 1.5kbp [11] and operons that crossed the start/end boundary of the contig or scaffold, to leave a final dataset of quality-checked *rrn* sequences. As previously reported [11], unlinked *rrn* operons were highly prevalent in the Deinococcus-Thermus and Planctomycetes phyla (Table S1). Each sequence in the final dataset was assigned a unique operon identifier and their nucleotide sequences were written to a single multifasta file using BEDTools [12].

To reduce computation time and aid analysis, a non-redundant (*NR 99.9%)* database was created for each dataset. First, a multifasta file of high-quality *rrn* operon sequences recovered from complete genomes was sorted by sequence length using BBTools sortbyname.sh [13] and clustered based on 99.9% nucleotide identity using vsearch -- cluster_smallmem [14]. First sorting by length ensured that the longest sequence in each cluster was retained as the representative sequence. The high-quality *rrn* operon sequences from incomplete assemblies were then appended to the representative sequence multifasta file and re-clustered, meaning that operon sequences from incomplete assemblies were only retained if they expanded on the sequence diversity of sequences from complete genomes. Consensus sequences for each NR cluster were generated using the --consout option.

We believe that the majority of users would benefit from using the NR 99.9% databases built from both complete and incomplete assemblies to maximise phylogenetic range while minimising computation burden. These are available for download as either representative (nrRep) or consensus (nrCon) sequences.

## DATABASE DESCRIPTION

The median length of *rrn* sequences was between 4,892 and 4,899 bp depending on database version (Table 1 & Figure 2A). ITS regions exhibited the greatest length variability across all database versions, followed by the 23S and 16S genes (Figure 2D-F). When considering complete genomes only, the corresponding source genomes have a mean per-genome rRNA copy number of 5.29 (*s* = 2.76) and 5.41 (*s* = 2.77) for the GTDB and RefSeq datasets, respectively (Figure 2B), while 63.62% or 63.80% of these genomes were represented in more than one NR cluster (Figure 2C), supporting previous reports of intragenomic diversity in rRNA genes and ITS regions [15–17]. For both datasets, approximately 81% of NR clusters contained only 1 *rrn* sequence (Figure 2G).

**Figure 2.**
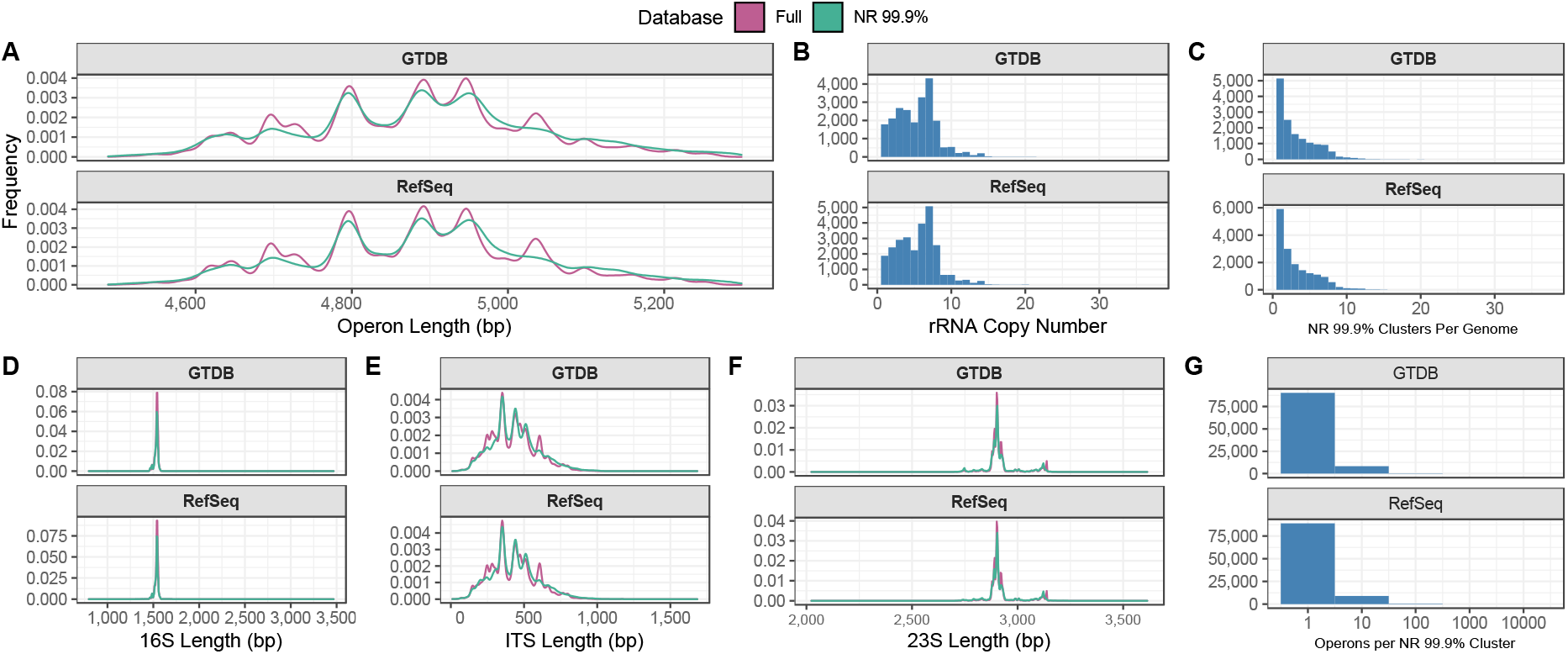
Summary statistics of the Full and non-redundant (NR 99.9%) FANGORN databases. A) length distribution of 16S-ITS-23S rRNA operons using the GTDB and RefSeq derived databases. B) number of distinct rRNA operons in genomes represented in the database. C) number of NR clusters per genome. D-F) length distribution of individual 16S-ITS-23S rRNA operon regions. G) number of rRNA operons represented by each NR cluster. Operons identified as outliers based on length (quartile 1/3 ± 1.5x interquartile range [IQR]) are not included in this plot to increase readability.

**Table 1.**
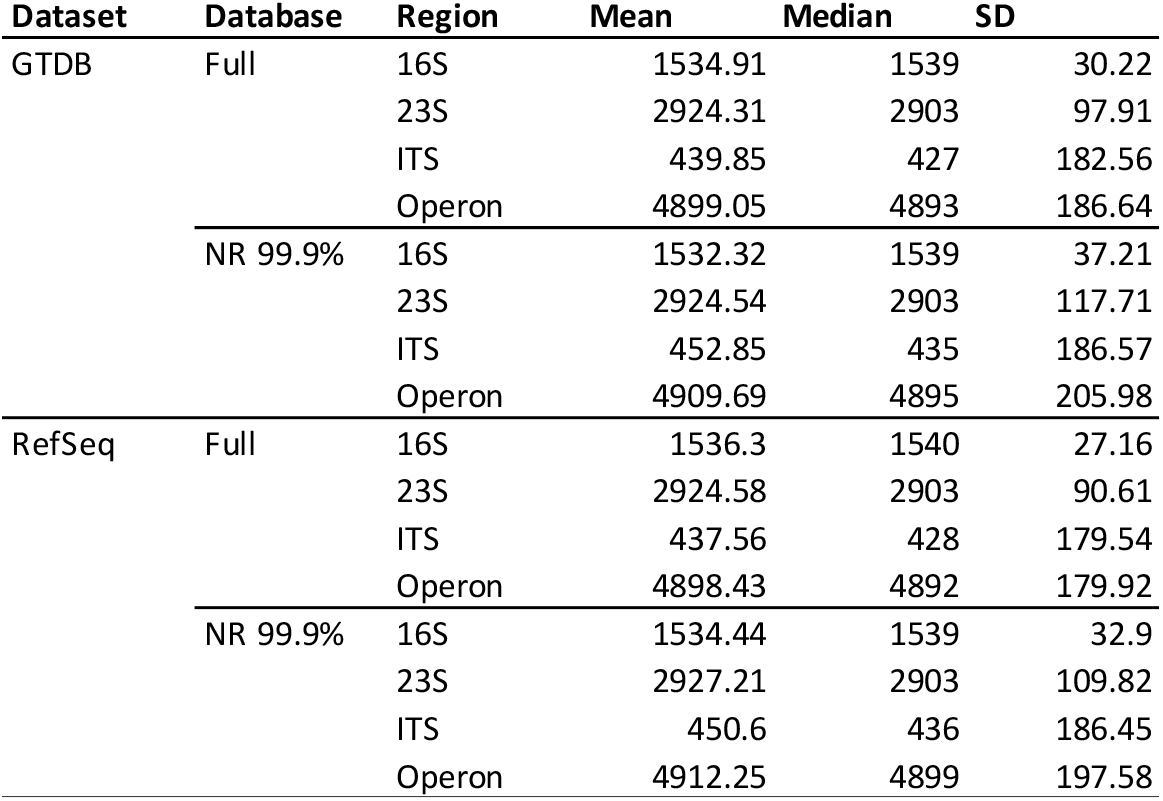
Summary statistics describing length (bp) of rrn operons, and constituent regions for each dataset and database.

**Table 2.**
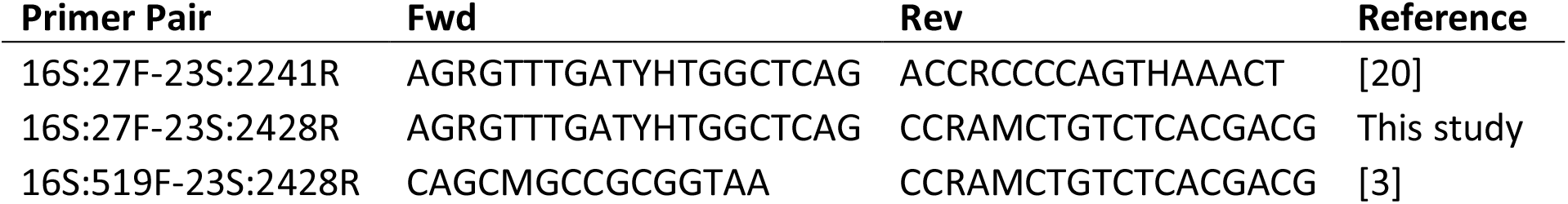
Primer sequences used for in silico PCR analysis.

## TAXONOMY

Taxonomy was assigned to each operon based on the source genome in seven-level format (Kingdom > Phylum > Class > Order > Family > Genus > Species). GTDB taxonomy was readily available to download in this format for archaea and bacteria. RefSeq TaxIDs were converted to this format by TaxonKit [18].

99.66% of GTDB NR clusters (and 97.18% of RefSeq clusters) exhibit 100% taxonomic agreement in the species-level taxonomy of their constituent operon sequences. In an effort to account for the absence of taxonomic agreement among the remaining clusters, three methods are used to assign taxonomy.

1. taxRep: source genome taxonomy of the cluster representative sequence
2. taxLCA: lowest common ancestor of all sequences in the cluster
3. taxMaj: lowest taxonomic rank at which there is a simple majority agreement of all sequences in the cluster

Files describing these taxonomy systems are available for download with the NR database. For most analyses, we recommend the taxRep system. The taxLCA and taxMaj systems are provided to compensate for clusters representing species with unclear or “disputed” taxonomy that may have arisen from incorrect taxonomic assignment of the genome when uploaded to the RefSeq database or an imperfect understanding of what defines a species. For example, a cluster which contains 97 sequences from *E. coli* and three sequences from the *Shigella* genus, two genera which are historically separated but should not be distinct based on genome-level comparative analysis [19], would be classified as s Escherichia coli by taxMaj, but as f_Enterobacteriaceae by taxLCA.

### PCR Primer Evaluation for Amplicon Generation

To assess the impact of primer choice on the functionality of the database, we tested three different primer combinations *in silico* (Table 1) using the perl script in_silico_pcr.pl (https://github.com/egonozer/in_silico_pcr) to evaluate their amplicon generation efficiency, phylogenetic bias, and amplicon length distribution. Two primer pairs from previous studies focused on the full-length rRNA operon were evaluated, in addition to a new pair that combines the forward and reverse primers from each pair to potentially generate a longer amplicon.

Based on the predicted primer binding characteristics there is a strong inverse relationship between predicted amplicon length and database coverage meaning there is a trade-off to be considered between phylogenetic range and resolution when selecting primers (Figure 4A & B). The primer pair 16S:27F-23S:2428R, which generates the largest amplicons, is predicted to be 1-2% less sensitive than the other two pairs (Figure 4A & Table S2).

**Figure 3.**
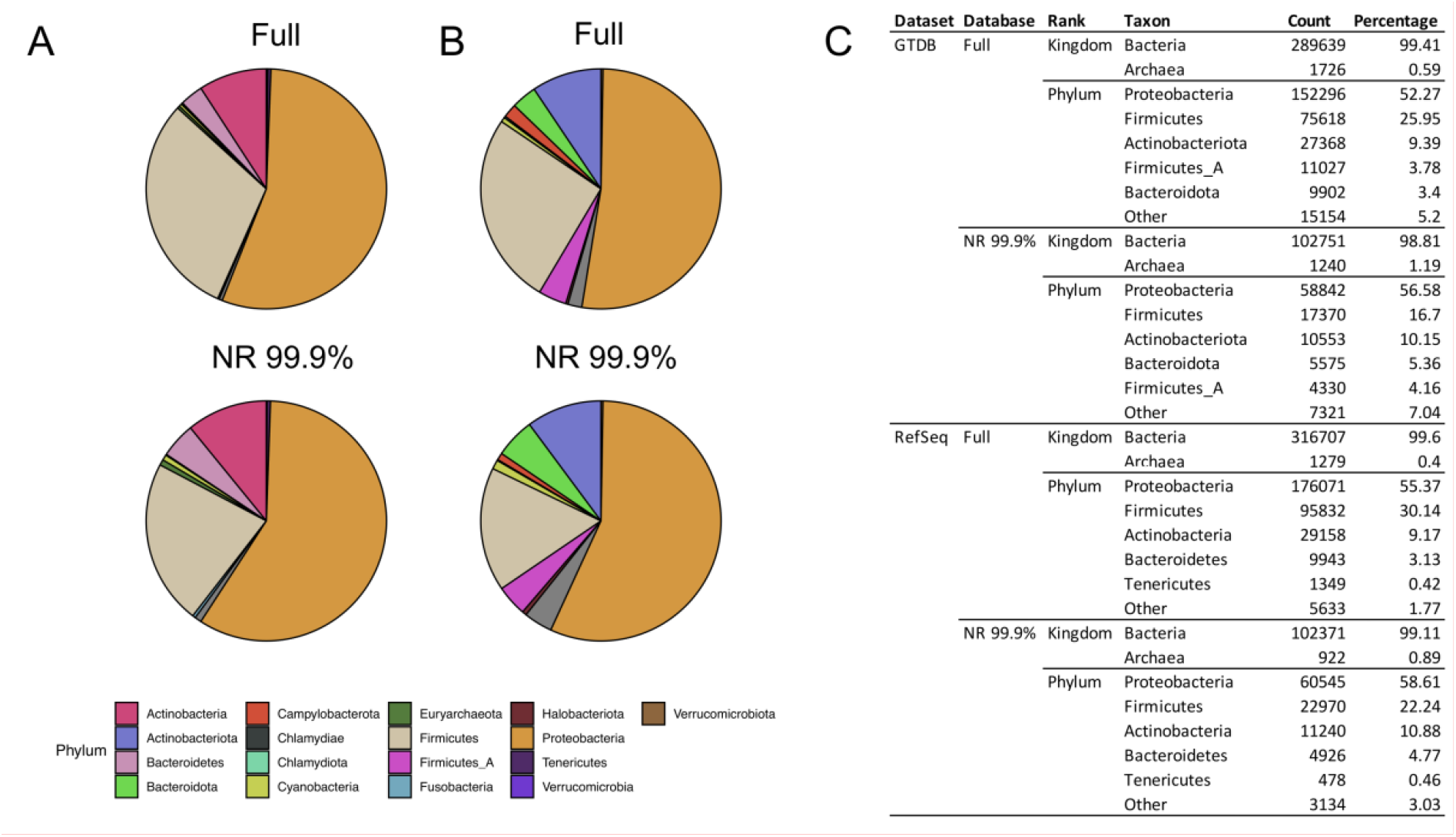
Taxonomic profile of FANGORN database. Phylum-level composition of Full and NR 99.9% databases using A) GTDB and B) RefSeq databases and taxonomy systems. C) Kingdom and Phylum-level composition expressed as number of operons and percentage of total.

**Figure 4.**
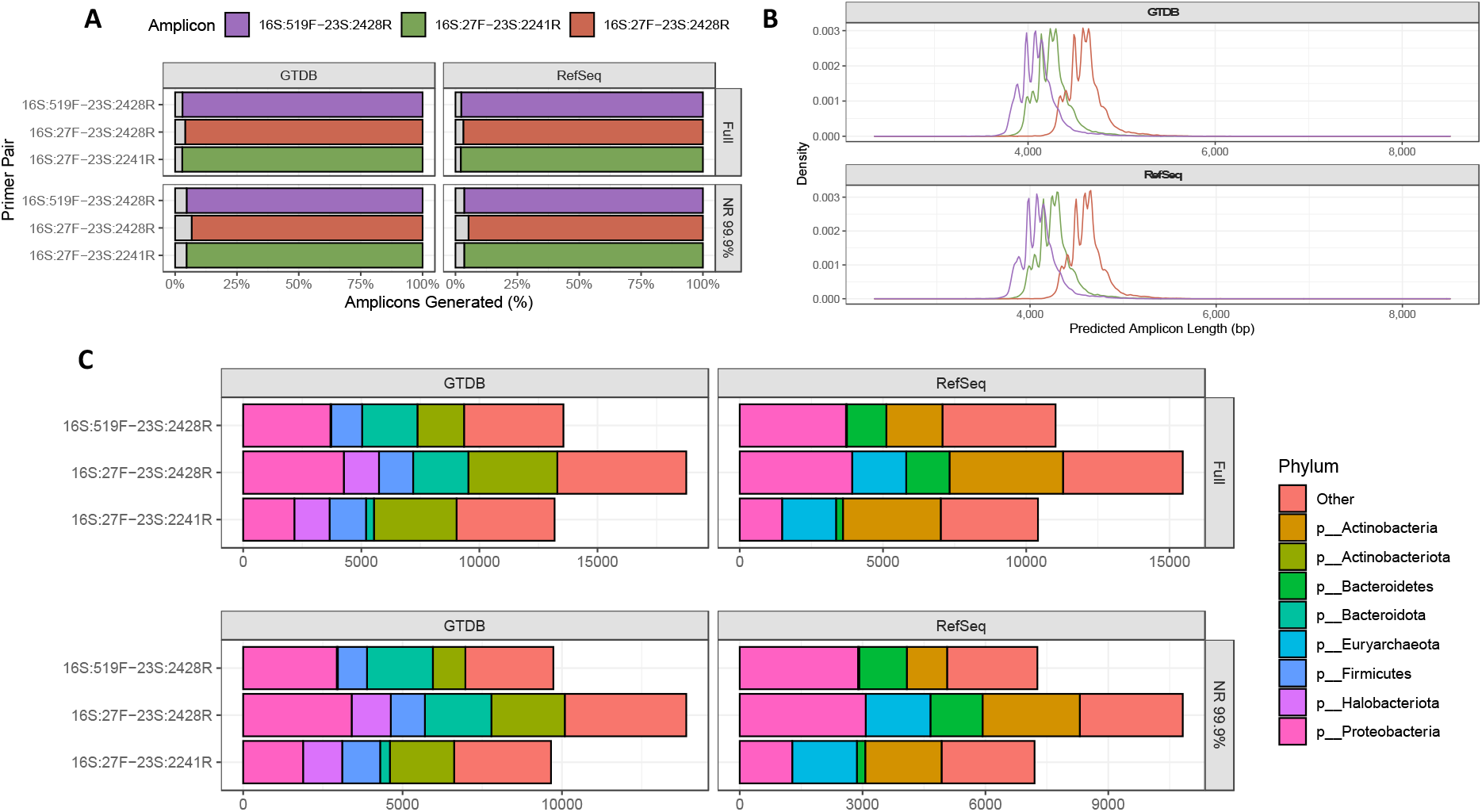
Predicted amplicon generation statistics. *A) Percentage of database sequences which were predicted to generate an amplicon by each primer pair. B) Predicted length distribution of amplicons generated from the* NR 99.9% *FANGORN database by each primer pair. C) Phylum-level composition of FANGORN database sequences where an amplicon was not predicted to be generated by each primer pair.*

Primer binding biases were predicted to be relatively consistent between primer pairs at phylum level (Figure 4C).

### Outlook

We predict that the future of rRNA-based phylogenetic analysis is linked to long-read sequencing technologies. As these become more reliable and scalable, we will begin to see more widespread adoption to exploit the more complete and detailed phylogenetic information contained in *rrn* operon sequences. Recent advances in long-read sequencing quality, such as UMI-tagging of template molecules [1], PacBio HiFi reads, and Oxford Nanopore Q20+ chemistry, mean that accessing this information is more easily achievable than ever. This will undoubtedly lead to a more detailed understanding of microbial diversity if the scientific community can act early to adopt standardised methods such as those currently recommended by the Earth Microbiome Project [21].

The FANGORN database will be updated with each major RefSeq and GTDB release to keep pace with the ever-growing collection of genome sequences and constantly evolving taxonomy systems.

## Data Availability

The FANGORN databases, and statistics describing genome length and *rrn* copy number for each taxon in the databases, are available to download from FigShare (doi: https://doi.org/10.26188/20086916). The code used to generate the databases is available at https://github.com/cazzlewazzle89/FANGORN.

## Supplementary Material

**Table S1.** Phylum-level distribution of genomes containing at least one unlinked rrn sequence. RefSeq genome assemblies and taxonomy labels are used and only phyla with> 10 genomes are shown.

| Phylum | # Genomes | % Genomes Containing Unlinked *rrn* Operons |
| --- | --- | --- |
| Deinococcus-Thermus | 73 | 76.71 |
| Candidatus Thermoplasmatota | 14 | 50 |
| Planctomycetes | 57 | 42.11 |
| Spirochaetes | 234 | 33.33 |
| Proteobacteria | 16221 | 2.82 |
| Crenarchaeota | 95 | 2.11 |
| Firmicutes | 6236 | 0.55 |
| Bacteroidetes | 1026 | 0.49 |
| Actinobacteria | 2689 | 0.26 |
| Tenericutes | 452 | 0.22 |

**Table S2.** In silico prediction of primer sensitivity. Sensitivity is expressed as the percentage of sequences in the database predicted to generate an amplicon using each primer pair.

| Dataset | Database | Primer Pair | Sensitivity (%) |
| --- | --- | --- | --- |
| GTDB | Full | 16S:27F-23S:2241R | 97.13417878 |
|  |  | 16S:27F-23S:2428R | 95.94872411 |
|  |  | 16S:519F-23S:2428R | 97.01714345 |
|  | NR 99.9% | 16S:27F-23S:2241R | 95.35632891 |
|  |  | 16S:27F-23S:2428R | 93.31769095 |
|  |  | 16S:519F-23S:2428R | 95.32171053 |
| RefSeq | Full | 16S:27F-23S:2241R | 97.85839628 |
|  |  | 16S:27F-23S:2428R | 96.83854006 |
|  |  | 16S:519F-23S:2428R | 97.67725623 |
|  | NR 99.9% | 16S:27F-23S:2241R | 96.51476867 |
|  |  | 16S:27F-23S:2428R | 94.7624718 |
|  |  | 16S:519F-23S:2428R | 96.48185259 |

